# Non-additivity of the functional properties of individual P450 species and its manifestation in the effects of alcohol consumption on the metabolism of ketamine and amitriptyline

**DOI:** 10.1101/2024.06.14.599105

**Authors:** Kannapiran Ponraj, Kari A. Gaither, Dilip Kumar Singh, Nadezhda Davydova, Mengqi Zhao, Shaman Luo, Philip Lazarus, Bhagwat Prasad, Dmitri R. Davydov

## Abstract

To explore functional interconnections between multiple P450 enzymes and their manifestation in alcohol-induced changes in drug metabolism, we implemented a high-throughput study of correlations between the composition of the P450 pool and the substrate saturation profiles (SSP) of amitriptyline and ketamine in a series of 23 individual human liver microsomes preparations from donors with a known history of alcohol consumption. The SSPs were approximated with linear combinations of three Michaelis-Menten equations with globally optimized *K*_M_ (substrate affinity) values. This analysis revealed a strong correlation between the rate of ketamine metabolism and alcohol exposure. For both substrates, alcohol consumption caused a significant increase in the role of the low-affinity enzymes. The amplitudes of the kinetic components and the total rate were further analyzed for correlations with the abundance of 11 major P450 enzymes assessed by global proteomics. The maximal rate of metabolism of both substrates correlated with the abundance of CYP3A4, their predicted principal metabolizer. However, except for CYP2D6 and CYP2E1, responsible for the low-affinity metabolism of ketamine and amitriptyline, respectively, none of the other potent metabolizers of the drugs revealed a positive correlation. Instead, in the case of ketamine, we observed negative correlations with the abundances of CYP1A2, CYP2C9, and CYP3A5. For amitriptyline, the data suggest inhibitory effects of CYP1A2 and CYP2A6. Our results demonstrate the importance of functional interactions between multiple P450 species and their decisive role in the effects of alcohol exposure on drug metabolism.

## l. Introduction

The functional properties of the human drug-metabolizing system are primarily determined by the properties of the ensemble of microsomal cytochromes P450 (P450) and their redox partners (microsomal monooxygenase, MMO) present in the endoplasmic reticulum (ER) of the cells of liver, intestine, lung, and most other tissues. It plays a key role in human drug metabolism and has become the major subject of studies on drug metabolism and disposition, drug-drug interaction, and adverse drug effects.

It has been long recognized that the wide inter-individual, age-dependent, and temporal variations in drug metabolism in the human population are dictated mainly by the respective changes in the composition of the P450 pool [1]. However, despite half a century of pharmacogenetic research, much of the interindividual variation in drug metabolism remains unexplained [2-3]. An important hurdle in this way is a lack of a comprehensive concept of the functioning of MMO as an integral multienzyme system, where multiple P450s compete for NADP-cytochrome P450 reductase (CPR) and interact with each other. There is growing evidence of a critical role of the functional effects of interactions between multiple P450 species [4-5]. Due to these largely unexplored effects, any change in the content of a particular P450 enzyme may affect drug metabolism in a complex, hard-to-predict manner.

Of particular importance is the functional impact of the changes induced by alcohol consumption. There are multiple indications of considerable modification of the pharmacokinetic properties of drugs by chronic alcohol exposure [6-10]. This study explores alcohol-induced changes in the functional properties of the cytochrome P450 ensemble using a collection of 23 samples of human liver microsomes (HLM) from donors with different degrees of alcohol exposure. As probe P450 substrates, we employed amitriptyline and ketamine, drugs used as antidepressants in patients with alcohol withdrawal syndrome [11-14] and metabolized by a wide spectrum of P450 species. We analyzed the variations in substrate saturation profiles (SSP) of these drugs and their correlation with the differences in the composition of the P450 ensemble characterized by a toolset of global proteomics.

Another objective of the study was to probe the additivity of the functional properties of the individual enzymes for drugs metabolized by multiple P450 species. A common and widely used approach for predicting the pathways of drug metabolism by P450 ensemble is based on a proportional projection (relative activity or total normalized rate) strategy [15-19], where the catalytic properties of all P450 enzymes metabolizing a given drug are presumed to be additive. Thus, the total rate of metabolism of a substrate is calculated as a sum of activities of all enzymes metabolizing it prorated for their fractional content in the P450 pool. In these calculations, researchers use the functional parameters of the individual P450 enzymes taken in isolation in reconstituted or genetically engineered systems. To improve the quality of prediction, semi-empirical scaling coefficients for *in vitro* to *in vivo* extrapolation are often used [19-21]. However, there is an increasing number of indications of wild oversimplification in this straightforward strategy due to ample mutual functional effects of multiple P450 enzymes in their ensemble. This study probes the applicability of the proportional projection approach in an attempt to reveal possible instances of non-additivity in the functional properties of the individual P450 species.

Understanding the functioning of MMO as an integral system requires advanced methods of analysis applied to the whole ensemble in its native environment in human liver microsomes (HLM). Combining advanced kinetic analysis with the toolkit of global proteomics provides a potent approach to unraveling complex functional interconnections in the P450 ensemble and elaborating new predictive techniques overcoming the limitations of the proportional projection strategy. In a pilot study with a fluorogenic substrate Coumarin 152 (C152) and a series of pooled HLM preparations, we introduced a strategy based on combining proteomic profiling of drug-metabolizing enzymes with global analysis of substrate saturation profiles (SSP) of polyspecific drug substrates [22]. Here we elaborated a new, improved version of our initial method where the PCA transform is used for a global fitting of a set of SSPs with a linear combination of several Michaelis-Menten equations with globally optimized *K*_m_ values.

This technique was used to probe the effect of alcohol exposure on the metabolism of ketamine and amitriptyline and explore the correlations between the composition of the P450 ensemble in HLM and the kinetic parameters of their P450-dependent demethylation. Our results reveal a considerable impact of alcohol exposure on the metabolism of the probe drugs and demonstrate the importance of considering the mutual functional effects of multiple P450 species for determining the pathways of drug metabolism.

## 2. Materials and Methods

### 2.1 Chemicals

Acetoacetanilide was procured from Tokyo Chemical Industry (Tokyo, Japan). S-ketamine and amitriptyline were the products of Cayman Chemical Company (Ann Arbor, MI, USA). S-Ketamine was obtained as a certified reference material (CRM) solution in methanol (1 mg/mL). The methanol solvent was evaporated under a stream of argon gas, and the dried substance was dissolved in 0.1 M Hepes buffer pH 7.4 containing 60 mM KCl. The concentration of ketamine in the stock solution was determined from its absorbance at 265 nm using the extinction coefficient of 0.56 mM^-1^cm^-1^. Amitriptyline was a product of Cayman Chemical Company (Ann Arbor, MI). Glucose-6-phosphate dehydrogenase from *Leuconostoc mesenteroides*, NADP, and Glucose-6-phosphate were the products of MilliporeSigma (Burlington, MA, USA). CYP3cide was a generous gift of Michael Wester (Vertex Pharmaceuticals, San Diego, CA, USA), All other reagents were of ACS grade and used without additional purification.

Pierce™ Trypsin Protease, MS Grade was purchased from Thermo Fisher Scientific. Stable isotope-labeled (heavy) peptides and synthetic unlabeled (light) peptides for targeted proteomics assay were purchased from Thermo Fisher Scientific (Rockford, IL) and New England Peptides (Boston, MA), respectively. All other reagents were of ACS grade and used without additional purification.

### 2.2 Pooled human liver microsomes

In this study, we used preparations of pooled human liver microsomes (HLM) from 50, 150, or 200 donors (mixed gender). The preparations CDN, DNJ, and WGP were INVITROCYP 150-Donor Human Liver Microsomes obtained from BioIVT Corporation (Westbury, NY). The lots EGW and ODN are INVITROCYP M-Class 50-Donor HLM procured from the same company. The lots 1910096 and 1210347 were XTreme 200 Mixed Gender HLM (200 donors) purchased from XenoTech LLC (Kansas City, KS), currently a part of BioIVT. The lot 2110263 corresponds to Mixed Gender HLM from 50 donors obtained from the same company. Lot 452116 of pooled mixed-gender HLM from 150 donors was procured from Corning Life Sciences Inc. (Tewksbury, MA). The suppliers of the HLM preparations used in our studies, BioIVT Corporation and Corning Inc. Life Sciences, have declared to adhere to the regulations of the Department of Health and Human Services for the protection of human subjects (45 CFR §46.116 and §46.117) and Good Clinical Practice (GLP), (ICH E6) in obtaining the samples of human tissues used for producing the preparations of human subcellular fractions available from these companies.

### 2.3 HLM samples from individual donors

In this study, we used 23 microsomal preparations from liver specimens obtained from individual donors. The set comprises 17 individual liver samples from the biobank established in the Prasad laboratory at WSU Spokane and 6 samples from donors with a history of moderate-to-heavy alcohol consumption procured from BioIVT Corporation (Baltimore, MD). The liver samples were included based on the availability of documented alcohol intake history. We identified and included donors with varying levels of alcohol consumption while excluding the use of illicit substances. Demographic characteristics of the donors may be found in the dataset available at Mendeley Data [23].

Preparation of microsomal fraction was performed as described by Nelson et al. [24] with minor modifications. Frozen liver tissue specimens (0.5 - 1 g) were covered with 0.1 M potassium phosphate buffer, pH 7.4, containing 0.125 M KCl, 0.25 M sucrose, and 1.0 mM EDTA (Buffer A) at room temperature and allowed to thaw. After decanting the buffer, the tissue was minced with scissors on ice and supplemented with 2.5 volumes of ice-cold buffer A containing 0.25 mM PMSF. The mixture was homogenized on ice with 10 strokes in a glass homogenizer with a motorized Teflon pestle. After diluting the homogenate to 7-8 volumes of the sample weight with ice-cold PMSF-containing buffer A, it was centrifuged at 10,500g for 40 min. The pellet was discarded, and the supernatant was centrifuged at 118,000 g for 90 min. The upper lipid layer and the supernatant were discarded. The pellet was resuspended in 0.1 M Na-HEPES buffer containing 150 mM KCl and 0.25 M sucrose, pH 7.4 (Buffer B) added to reach 1.5 - 2 volumes of the sample weight and centrifuged at 118,000 g for 90 min. The pellet was resuspended in Buffer B (0.5 ml/g of tissue) with a syringe and plastic pestle in a 1.5 ml microfuge tube and stored at -80 °C.2.4. Microsomes containing recombinant human cytochromes P450

Most of the preparations of insect-cell microsomal containing baculovirus-expressed individual P450 enzymes (Supersomes^TM^) were the products of Gentest (Huntsville, AL), now, a part of Discovery Life Sciences (Woburn, MA). In the present study, we used the preparations containing CYP2B6 (lot 31487), CYP2C8 (lot 81760), CYP2C9 (lot 75854), CYP2C19 (lot 73445), CYP2E1 (lot 23012), CYP3A4 (lot 81745), CYP3A5 (lot 89573) and CYP4A11 (lot 2402280). All those preparations contained human CPR and cytochrome *b*_5_ co-expressed. The preparations of insect cell microsomes containing human CYP2A6 (lot 2520331) and CYP2D6 (lot 2350106) along with human CPR and cytochrome b_5_ (Baculosomes®) were procured from Thermo Fisher Scientific (Waltham, MA, USA).

### 2.5 Proteomic characterization of P450 abundances in HLM

The pooled HLM preparations were digested using trypsin using an optimized protocol [25]. The abundances of 11 major P450 species, cytochrome *b*_5_, and CPR were quantified by a targeted proteomics assay as described earlier [23, 26]. Relative abundance of P450 species and protein markers of alcohol exposure (HSPA5, PDIA3, P4HB, and CES2 [27]) in the individual and pooled HLM preparations were determined based on the global proteomics-based total protein approach as described in [25].

### 2.6 Genotypic characterization of individual HLM samples

Human liver specimens were minced to extract genomic DNAs (gDNAs) using the PureLink™ Genomic DNA Mini Kit (Invitrogen™, K182001), and genomic DNA concentrations were measured using a NanoDrop™ 2000/2000c spectrophotometer (Thermo Scientific™, USA). The CYP2D6*2 (2851 C>T, rs16947), CYP2D6*10 (100 C>T, rs1065852), CYP2B6*5 (25505 C>T, rs3211371), CYP2B6*4 (18053 A>G, rs2279343) and CYP2B6*6 (15631 G>T, rs3745274; 18053 A>G, rs2279343) SNPs were detected by real-time PCR using 10-20 ng of genomic DNA in a 5-uL PCR reaction system containing TaqMan™ Genotyping Master Mix (Applied Biosystems™, 4371355) and either the TaqMan™ SNP Genotyping Assay (Applied Biosystems™, 4351379) or the TaqMan™ Drug Metabolism Genotyping Assay (Applied Biosystems™, 4362691) according to the manufacturer’s user guide. All real-time PCR was performed in the Washington State University-Spokane Genomics Core Facility using a Bio-Rad CFX384 real-time PCR machine with Bio-Rad CFX Manager^TM^ 3.1 to characterize genotypes. Four replicates were conducted for each sample in each experiment.

### 2.7 Characterization of the content of protein, NADPH-cytochrome P450 reductase, and cytochromes P450 and b_5_ in HLM

Protein concentrations in microsomal suspensions were determined by the bicinchoninic acid assay [28]. The concentration of CPR in microsomal membranes was determined based on the rate of NADPH-dependent reduction of cytochrome *c* at 25 °C, and the effective molar concentration of CPR was estimated using the turnover number of 3750 min^-1^ [29]. The total concentration of cytochromes P450 in HLM was determined with a variant of the “oxidized CO versus reduced CO difference spectrum” method described earlier [29].

### 2.7 High-throughput assays of oxidative demethylation of ketamine and amitriptyline

Monitoring of ketamine and amitriptyline demethylation was performed with a high-throughput fluorimetric assay of formed formaldehyde based on the Hantzsch reaction with acetoacetanilide [30]. A series of substrate saturation profiles (SSP) were obtained with ketamine and amitriptyline concentrations decreasing from 700 μM (ketamine) or 734 μM (amitriptyline) with a dilution factor of 1.73333 (z✓3). Reactions were carried out at 30 °C in 0.1 M Na-Hepes buffer, pH 7.4, containing 60 mM KCl, 200 μM NADP, 2 mM glucose-6-phosphate and 1 U/ml of glucose-6-phosphate dehydrogenase. The concentration of HLM protein in the incubation media was 0.4-0.8 mg/ml. Incubation times for the ketamine and amitriptyline assays were 20 and 30 min., respectively.

Automation of the assay was achieved by using an OT-2 liquid handling robot (Opentrons Inc., Brooklyn, NY, USA). The spectra of excitation in the 300 - 430 nm region (emission at 468 nm) were recorded using a Cary Eclipse fluorometer equipped with a plate reader accessory (Agilent Technologies, Santa Clara, CA, USA). A detailed description of the assay may be found in our earlier publication [30]. To obtain a dataset of SSPs for global analysis, we averaged from 4 to 7 individual SSPs obtained with each HLM preparation after their polynomial smoothing with a 3-point window in semilogarithmic (log(X)) coordinates.

### 2.8 Enzyme inhibition experiments

Inhibition of ketamine and amitriptyline metabolism by CYP3cide was studied using the high-throughput assay described above. The substrate concentration was kept constant at 140 μM and 250 μM for ketamine and amitriptyline, respectively. The concentration of CYP3cide in the series of 12 samples in each assay decreased from 6.4 μM (ketamine) or 3.2 μM (amitriptyline) to zero with a dilution factor of 1.414 (≈√2). The fractional inhibition at each inhibitor concentration was determined from the ratio of the reaction rate in the presence of an inhibitor to the rate observed at no inhibitor added. The resulting dependence of the fractional inhibition on the inhibitor concentration was fitted to a hyperbolic equation [31]:

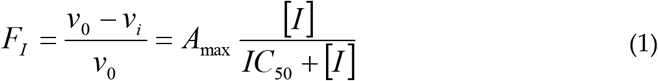

In this equation, *F*_I_ and *v*_i_ are the fractional inhibition and the reaction velocity observed at inhibitor concentration [I], *v*_0_ is the reaction rate at no inhibitor added, and *A*_max_ is the maximal amplitude of inhibition. *IC*_50_ is the concentration of inhibitor at which the inhibition reaches 50% of its maximal amplitude (*A*_max_).

### 2.9 Global kinetic analysis of substrate saturation profiles

The datasets for global kinetic analysis were assembled as the sets of averages of 4 - 7 SSP obtained with each HLM preparation. The datasets obtained with pooled and individual HLM samples, either separately or in combination, were subjected to Principal Component Analysis. The results of PCA of the pooled HLM dataset were used to find a set of three Michaelis-Menten equations whose linear combinations best approximate each of the individual SSP traces. To find the *K*_m_ values of these three Michaelis-Menten components we used a Neilder-Mead optimization procedure combined with the SURFIT algorithm of multidimensional linear regression [32] applied to a set of three first principal component vectors and the average of all traces in the dataset (the base vector of the analysis). The optimized set of three Michaelis-Menten components found thereby was used to resolve the individual kinetic phases (high-, medium-, and low-affinity) in each of the SSPs in pooled, individual, or combined SSPs datasets using the SURFIT algorithm [32]. In the cases where the SURFIT algorithm yielded a negative amplitude for one of the three Michaelis-Menten components, it was excluded from the fitting, and the approximation was done with the two remaining components. The square correlation coefficients for these approximations were always >0.99 and, in most cases, >0.995.

To further improve the results of global analysis and eliminate possible bias caused by individual fitting of the SSPs, we then approximated the obtained profiles of *V*_max_ values by linear combinations of vectors of eigenvalues of the five first principal components. The resulting approximations were then used to analyze the correlations between the shapes and the amplitudes of SSPs with the composition of the cytochrome P450 ensemble in HLM samples.

All manipulations with the dataset, Principal Component Analysis, and regression analysis were performed using our SpectraLab software [4, 33], which is available for download at http://cyp3a4.chem.wsu.edu/spectralab.html.

### 2.10 Analysis of correlations between the SSPs and the composition of the P450 pool

To find the correlation of the shape and amplitudes of SSP with variations of the abundances of individual P450 enzymes, we calculated the vectors of relative fractional abundance (VFRA) for 11 major P450 in HLMs (CYP1A2, CYP2A6, CYP2B6, CYP2C8, CYP2C9, CYP2C19, CYP2D6, CYP2E1, CYP3A4, CYP3A5, and CYP4A11). For these calculations, the raw MS intensities (MSI) for each (*i*-th) P450 protein in each (*j-*th) HLM sample in the dataset of 23 individual HLM samples or the combined data set of 23 individual and 9 pooled preparations were first normalized on the sum of intensities of 75 drug-metabolizing enzymes (DME) and ER-stress-related proteins with known localization in the microsomal membrane or microsomal lumen (see [27] for the list of these proteins):

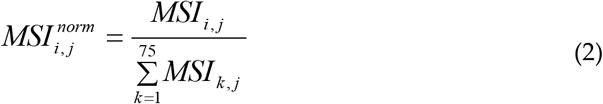

The purpose of this normalization is to account for possible contamination of the HLM samples by non-microsomal proteins. We also calculated the total normalized MSI of P450 enzymes as a sum of all MSI values of P450 enzymes in the set under analysis 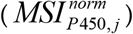.

After normalization, the averaged normalized intensity for each P450 protein in all HLM in the set was assessed and used to calculate the VRA value for each (*i*-th) protein in each (*j*-th) HLM sample:

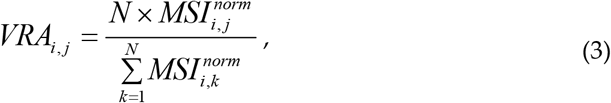

where N is the total number of HLM preparations in the dataset (23 for the set of individual HLMs or 32 for the combined set of pooled and individual samples).

To obtain the vector of total P450 abundance in the samples we used the results of CO-differential spectroscopy. The total concentrations of P450 proteins in all HLM samples determined with a variant of the “oxidized CO versus reduced CO difference spectrum” [29] were normalized on their average. The resulting vector of the relative total abundance of P450 proteins was then used to calculate the relative fractional contents of each (*i*-th) protein in each (*j*-th) HLM sample yielding the vector of relative fractional content (VFC) of P450 species:

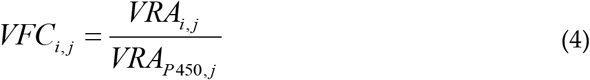

Analysis of correlations between the composition of the P450 ensemble and the shape and amplitude of SSP was based on approximating the profiles of the amplitudes of each Michaelis-Menten component (*V*_max1_, *V*_max2_, and *V*_max3_) and their total (*V*_max total_) with linear combinations of the vectors of relative fractional content of eleven major cytochrome P450 species. For these trials, the vectors of the *V*_max_ values for the HLM samples in the dataset were fitted to linear combinations of one to four vectors of fractional content of P450 species using the SURFIT algorithm of multidimensional linear regression [32]. We wrote a script for the SpectraLab software which successively applies the SURFIT algorithm to all possible combinations of up to four vectors of relative abundances of the P450 species under analysis and identifies the best approximation (in terms of the square correlation coefficient).

## 3. Results

### 3.1 Metabolism of ketamine and amitriptyline by the individual P450 species

Although the role of the individual P450 enzymes in the demethylation of ketamine [34-37] and amitriptyline [38-41] has been already addressed in the literature, our knowledge of the catalytic parameters of drug-metabolizing P450 species with these substrates remains essentially incomplete. In this study, we used our high-throughput technique to examine the kinetics of P450-catalyzed demethylation reactions [30] to characterize the ability of eleven human cytochrome P450 species (1A2, 2A6, 2B6, 2C8, 2C9, 2C19, 2D6, 2E1, 3A4, 3A5, and 4A11) to metabolize ketamine and amitriptyline. Results of these assays are shown in Tables 1 and 2, where the P450 species are sorted in the order of their decreasing abundance in HLM.

**Table 1.** Parameters of S-Ketamine metabolism by recombinant P450 enzymes_*_.

| P450 species | Averaged fraction in the P450 pool, % | $K_M$, $\mu\text{M}$ | $V_{\max}$, $\text{min}^{-1}$ | $\text{Cl}_{\text{int}}$, $\mu\text{M}^{-1} \text{min}^{-1}$ | % of turnover | | |
| --- | --- | --- | --- | --- | --- | --- | --- |
| | | | | | at 50 $\mu\text{M}$ | at 250 $\mu\text{M}$ | at saturation |
| CYP3A4 | 31.0 ± 2.1 | 113 ± 18 | 44.0 ± 7.7 | 0.390 | 69 | 70 | 64 |
| CYP2C9 | 19.3 ± 0.5 | 250 ± 76 | 6.2 ± 2 | 0.025 | 3.3 | 4.4 | 5.6 |
| CYP2E1 | 17.0 ± 1.4 | 457 ± 182 | 5.3 ± 2.1 | 0.012 | 1.5 | 2.3 | 4.2 |
| CYP2A6 | 11.1 ± 0.4 | 79 ± 30 | 5.6 ± 1.6 | 0.071 | 4.0 | 3.5 | 2.9 |
| CYP1A2 | 10.9 ± 0.9 | 236 ± 48 | 9.5 ± 0.6 | 0.040 | 3.0 | 4.0 | 4.9 |
| CYP2D6 | 2.7 ± 0.2 | 685 ± 124 | 68.0 ± 24.3 | 0.099 | 2.1 | 3.6 | 8.5 |
| CYP2C8 | 2.7 ± 0.1 | 152 ± 49 | 5.6 ± 0.4 | 0.037 | 0.6 | 0.69 | 0.70 |
| CYP2B6 | 2.6 ± 0.6 | 54 ± 14 | 50.6 ± 8 | 0.938 | 11 | 8.1 | 6.2 |
| CYP3A5 | 1.1 ± 0.2 | 103 ± 22 | 27.5 ± 3.7 | 0.267 | 1.6 | 1.5 | 1.4 |
| CYP2C19 | 1.0 ± 0.1 | 19 ± 7 | 13.9 ± 7.1 | 0.719 | 1.6 | 0.94 | 0.64 |
| CYP4A11 | 0.7 ± 0.1 | 28 ± 7 | 29.8 ± 5.5 | 1.080 | 2.2 | 1.4 | 0.98 |

<sup>a</sup> Averaged fractional content of P450 species obtained in the analysis of 12 pulled HLM preparations by MRM proteomics (see dataset available at Mendeley Data [23]).
<sup>\*</sup> The values of the parameters obtained from our experiments shown in this table represent the averages of the results of 4-7 individual experiments. The ± values correspond to the confidence intervals calculated for $p = 0.05$ . The $V_{\max}$ values are expressed as mols of formaldehyde formed per minute per mole of cytochrome P450 ( $\text{min}^{-1}$ ).

**Table 2.**
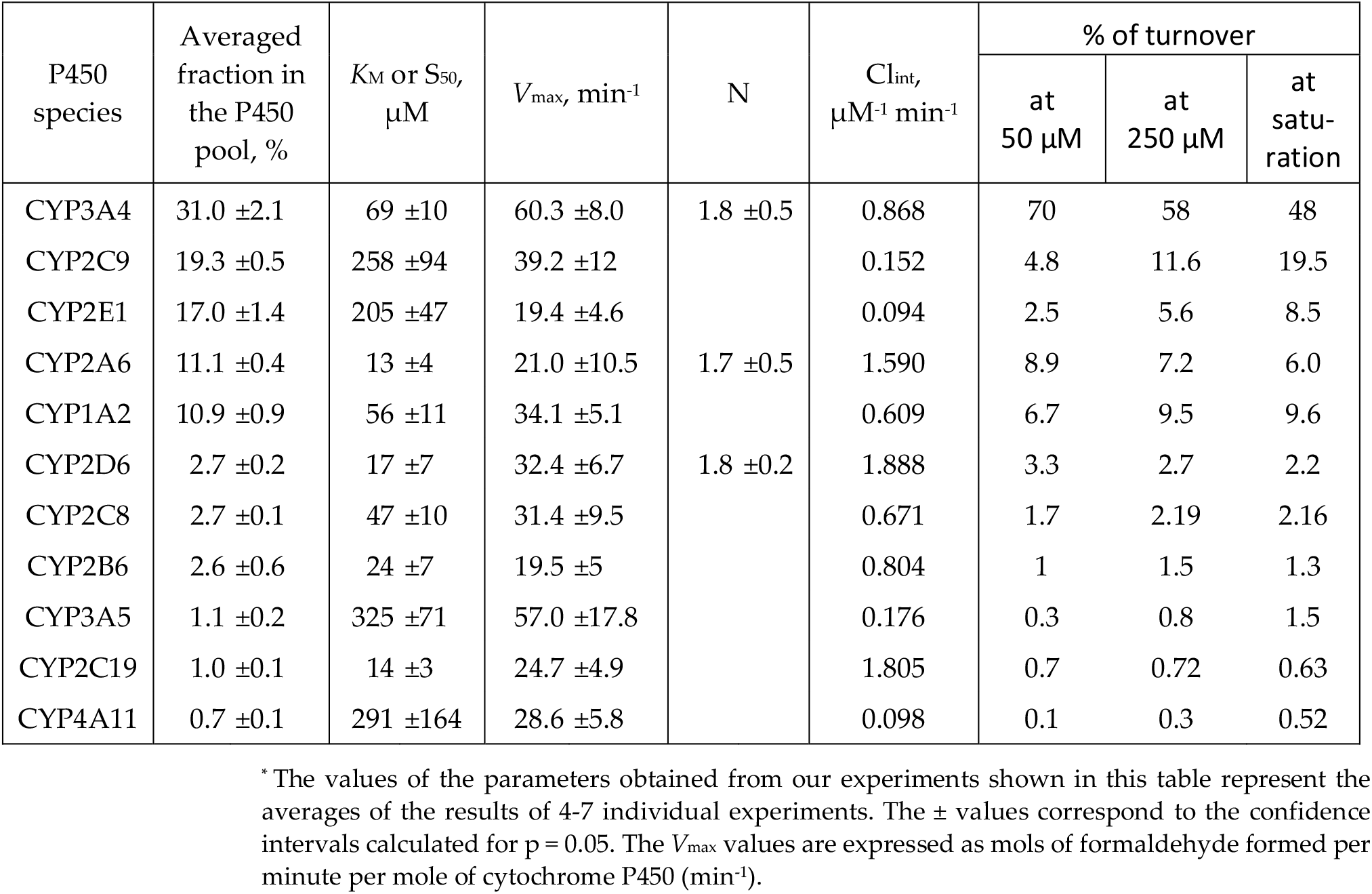
Parameters of amitriptyline metabolism by recombinant P450 enzymes_*_.

| P450 species | Averaged fraction in the P450 pool, % | $K_M$ or $S_{50}$, $\mu M$ | $V_{max}$, $\text{min}^{-1}$ | N | $Cl_{int}$, $\mu M^{-1} \text{min}^{-1}$ | % of turnover | | |
| --- | --- | --- | --- | --- | --- | --- | --- | --- |
| | | | | | | at 50 $\mu M$ | at 250 $\mu M$ | at saturation |
| CYP3A4 | 31.0 ± 2.1 | 69 ± 10 | 60.3 ± 8.0 | 1.8 ± 0.5 | 0.868 | 70 | 58 | 48 |
| CYP2C9 | 19.3 ± 0.5 | 258 ± 94 | 39.2 ± 12 | | 0.152 | 4.8 | 11.6 | 19.5 |
| CYP2E1 | 17.0 ± 1.4 | 205 ± 47 | 19.4 ± 4.6 | | 0.094 | 2.5 | 5.6 | 8.5 |
| CYP2A6 | 11.1 ± 0.4 | 13 ± 4 | 21.0 ± 10.5 | 1.7 ± 0.5 | 1.590 | 8.9 | 7.2 | 6.0 |
| CYP1A2 | 10.9 ± 0.9 | 56 ± 11 | 34.1 ± 5.1 | | 0.609 | 6.7 | 9.5 | 9.6 |
| CYP2D6 | 2.7 ± 0.2 | 17 ± 7 | 32.4 ± 6.7 | 1.8 ± 0.2 | 1.888 | 3.3 | 2.7 | 2.2 |
| CYP2C8 | 2.7 ± 0.1 | 47 ± 10 | 31.4 ± 9.5 | | 0.671 | 1.7 | 2.19 | 2.16 |
| CYP2B6 | 2.6 ± 0.6 | 24 ± 7 | 19.5 ± 5 | | 0.804 | 1 | 1.5 | 1.3 |
| CYP3A5 | 1.1 ± 0.2 | 325 ± 71 | 57.0 ± 17.8 | | 0.176 | 0.3 | 0.8 | 1.5 |
| CYP2C19 | 1.0 ± 0.1 | 14 ± 3 | 24.7 ± 4.9 | | 1.805 | 0.7 | 0.72 | 0.63 |
| CYP4A11 | 0.7 ± 0.1 | 291 ± 164 | 28.6 ± 5.8 | | 0.098 | 0.1 | 0.3 | 0.52 |

\* The values of the parameters obtained from our experiments shown in this table represent the averages of the results of 4-7 individual experiments. The ± values correspond to the confidence intervals calculated for $p = 0.05$ . The $V_{max}$ values are expressed as mols of formaldehyde formed per minute per mole of cytochrome P450 ( $\text{min}^{-1}$ ).

Although our results are consistent with the earlier studies pointing out CYP3A4, CYP2B6, and CYP2C9 as the major metabolizers of ketamine [35-37] and CYP3A4, CYP2C19, and CYP2D6 as the most efficient catalysts of amitriptyline demethylation [38, 40-41], we revealed some other, yet unidentified, important players in the metabolism of these substrates. Thus, surprisingly, CYP4A11 was identified as the most efficient metabolizer of ketamine. This enzyme is followed by CYP2B6, CYP2C19, and CYP3A4 (Table 1). Importantly, the highest *V*_max_ with ketamine is exhibited by CYP2D6, which, however, has a low affinity to this substrate (Table 1). In the case of amitriptyline, the most efficient metabolizers are CYP2D6, CYP2C19, CYP2A6, and CYP3A4 (Table 2). According to our results, in terms of catalytic efficiency CYP2D6 outperforms CYP2C19, which was assumed the most potent metabolizer of amitriptyline in the earlier studies [38, 40]. New in this list is CYP2A6, which has not been yet probed on its ability to metabolize amitriptyline.

Notably, the turnover rates for amitriptyline demethylation observed in our experiments (19 - 60 min^-1^, Table 2) are considerably higher than those reported in the literature (1.5 - 15 min^-1^ [38, 40-41]). This apparent discrepancy is likely caused by a difference in the methods of detecting the demethylation products. In our assays, we quantified the formation of formaldehyde produced in the demethylation of the dimethylamino group of the substrate. Thus, our results reflect the formation of both possible products nortriptyline (mono-desmethyl amitriptyline) and desmethyl nortriptyline. In contrast, all previous studies were based on detection of nortriptyline only. An apparent underestimation of the rate of amitriptyline demethylation in the earlier studies points out desmethyl nortriptyline as the predominant product of the metabolism of this substrate by the ensemble of human P450 enzymes.

Using the parameters of ketamine and amitriptyline metabolism by P450 isoenzymes, we estimated the fractional involvement of the individual P450 species in the N-demethylation of these substrates. In these calculations, we followed the “total nor-malized rate” (TNR) approach proposed by Rodrigues [15] and further developed by others [17-19]. This approach is based on a proportional projection of the presumed activities of the individual P450 enzymes to their ensemble with a known composition. The fractional content of 11 P450 species used in these calculations is based on averaging their abundances in 14 pulled HLM preparations determined by MRM) based targeted proteomics (see dataset available at Mendeley Data [23]). The results of these calculations are also shown in Tables 1 and 2.

As seen from these tables, for both substrates the predominant role in their metabolism is expected to be played by CYP3A4. The CYP3A4-dependent N-demethylation of ketamine is estimated to account for 70% of the total rate at 250μM substrate while the CYP2B6, CYP2C9, CYP1A2, and CYP2D6 enzymes are expected to take the responsibility of another 20% of the metabolism (Table 1). The leading role of CYP3A4, CYP2B6, CYP2C9, and CYP1A2 has already been predicted [34-37] while a contribution of CYP2D6 has not been yet reported. Having the highest turnover rate with ketamine, this enzyme may be responsible for up to 8.5% of its demethylation at saturating concentrations of the drug.

In the case of amitriptyline, CYP3A4 is presumed to take the responsibility of 58% of metabolism at 250 μM substrate concentration. It is followed by CYP2C9, CYP1A2, CYP2A6, and CYP2E1 whose cumulative involvement is expected to be around 34% (Table 2). These results do not corroborate the predominant role of CYP2C19 inferred in the previous publications [38, 40]. Although this enzyme has a high affinity to amitriptyline and a modest turnover rate (Table 2), its involvement is diminished by its low content in HLM. While the involvement of CYP3A4, CYP2C9, and CYP1A2 has already been demonstrated [40], a considerable potential role of CYP2A6 and CYP2E1 was unexpected. The predicted involvement of CYP2E1, which has a low affinity to amitriptyline, increases at high concentrations of the drug and comprises up to 8.5% at substrate saturation.

### 3.2 Assembling a set of individual donor-derived HLM preparations and characterizing genetic variability of cytochromes P450 across the donors

To explore the relationship between the degree of alcohol exposure, the composition of the P450 ensemble, and the parameters of ketamine and amitriptyline demethylation we constructed a set of SSPs in a series of 23 individual HLM preparations where the composition of the microsomal drug-metabolizing ensemble was characterized by global proteomics based on the total protein approach [25]. The HLM samples in this series were selected to represent different degrees of alcohol exposure of the liver donors. It contained 9 HLM samples from non-and light drinkers (<=1 drink per day), 7 samples from moderate drinkers (2-3 drinks per day), and 7 samples from heavy drinkers (>=4 drinks per day). Demographic characteristics of the donors and the results of proteomic characterization of HLM preparations may be found in the dataset available at Mendeley Data [23].

Prior to probing the correlations of the kinetics of ketamine and amitriptyline metabolism with the composition of the P450 pool in HLM, we sought to probe genetic polymorphism in the most genetically variable P450 enzymes in our set of individual HLMs. The presence of poor- or ultra-rapid genetic P450 variants in some samples of our set may affect the observed variability in drug metabolism.

When considering the potential effect of single-nucleotide polymorphisms (SNPs), we focused on the genetic variants known to affect the rates of enzyme turnover or substrate specificity. SNPs affecting the level of expression were excluded as the respective changes in the protein abundances are anyway reflected in the results of proteomic analysis. In selecting P450 enzymes to analyze for genetic variation, we focused on those possessing SNPs of interest with an overall frequency of 0.2 or more. When identifying SNPs to focus on, we set the occurrence threshold at 0.1. At these thresholds, we expected to find at least one homo- or heterozygote of the SNPs of interest in our set of 23 HLM samples.

The most genetically variable P450 enzymes among the principal metabolizers of ketamine and amitriptyline are CYP2D6, CYP2B6, and CYP2C9 [42-43]. Considering the genetic polymorphism in CYP2C9, we found two major SNPs affecting the enzyme activity: 3608C>T (rs1799853, R144C) and 42614A>C (rs1057910, I359L). However, according to the NCBI NLM database (https://www.ncbi.nlm.nih.gov/snp), their total frequency of occurrence is below 0.2. Thus, we refrained from analyzing CYP2C9 polymorphism in this study.

The most frequent functionally-important SNPs in CYP2D6 are 2851C>T (rs16947, R296C, CYP2D6*2) and 100C>T (rs1065852, P34S, CYP2D6*10) [42]. Both have high frequencies (0.3 - 0.4 and -0.2, respectively) and pronounced functional consequences (largely decreased turnover rate) [44-46]. All other SNPs were excluded due to a lack of demonstrated functional effect or low frequency.

In the case of CYP2B6 we focused on 18053A>G (rs2279343, K262R, CYP2D6*4 and CYP2D6*6), 25505C>T (rs3211371, R487C, CYP2D6*5), and 15631G>T (rs3745274, Q172H, CYP2D6*6) SNPs. Their total frequency is around 0.5 and the individual frequencies are around 0.15, 0.1, and 0.3, respectively. All three SNPs have significant functional effects - K262R considerably increases the turnover rate and changes substrate specificity [47-48], while the other two are detrimental to the enzyme activity [49]. All other CYP2B6 SNPs were considered out of interest.

The occurrences of the above-mentioned CYP2B6 and CYP2D6 SNPs were analyzed in 17 samples from our HLM set for which we had genetic material. The results of this analysis are shown in Table 3. By analyzing these results, we identified HLM samples that are homo- or heterozygous for CYP2B6 or CYP2D6 genetic variants of interest. SNP heterozygotes with major allelic variants (CYP2B6*1 or CYP2D6*1) were not considered because the functional properties of P450 enzymes are likely to be unaffected in this case. Selected HLM samples, where the function of CYP2B6 or CYP2D6 may be affected by the presence of SNPs, are highlighted in bold in Table 3. To probe the potential effects of genetic polymorphisms, we compared the results obtained with the full set of HLM samples with the results obtained upon omitting the samples listed in bold in Table 3 when analyzing the correlation of the kinetic parameters of ketamine and amitriptyline metabolism with the composition of the P450 pool.

**Table 3.** CYP2B6 and CYP2D6 genotypes in the individual HLM preparations from the biobank in Prasad’s laboratory.

| HLM sample ID<sup>a</sup> | CYP2D6 genotype | CYP2B6 genotype | HLM sample ID | CYP2D6 genotype | CYP2B6 genotype |
| --- | --- | --- | --- | --- | --- |
| **HMC591** | **\*2/\*2** | **\*1/\*1** | HFC583 | *1/*1 | *1/*6 |
| **HFAA510** | **\*2/\*10** | **\*1/\*6** | HMC593 | *1/*10 | *1/*6 |
| **HFH544** | **\*2/\*10** | **\*1/\*1** | HFC640 | *1/*2 | *1/*5 |
| **HMC714** | **\*2/\*10** | **\*1/\*6** | HMC671 | *1/*10 | *1/*5 |
| **HMC765** | **\*2/\*10** | **\*1/\*6** | HMC750 | *1/*10 | *1/*1 |
| **HMC820** | **\*2/\*10** | **\*1/\*6** | HFC791 | *1/*1 | *1/*6 |
| **HMC682** | **\*1/\*1** | **\*5/\*5** | HMC808 | *1/*1 | *1/*5 |
| **HMC529** | **\*1/\*10** | **\*5/\*6** | HMC846 | *1/*10 | *1/*5 |
| **HMC531** | **\*1/\*10** | **\*5/\*6** |  |  |  |

<sup>a</sup> The HLM identifiers of homo- and heterozygotes of minor CYP2B6 and CYP2D6 variants are highlighted in bold.

### 3.3 Metabolism of ketamine by HLM

A series of titration curves for ketamine demethylation obtained with a series of nine distinct pooled HLM preparations is shown in Fig. 1a. All shown SSPs are normalized by the concentration of CPR in HLM samples. As seen from this figure, the probed pooled HLM preparations differ considerably in both the shape and the amplitude of the kinetic traces. For quantitative characterization of this variability, we applied a global fitting approach based on the use of Principal Component Analysis (PCA). The main panel in Fig. 1b shows the first three principal components (PC) obtained by PCA applied to the dataset shown in Fig. 1a. Assuming that these PCs represent different combinations of one and the same set of several Michaelis-Menten dependencies, we used a combination of linear multidimensional regression algorithm with Neilder-Mead nonlinear regression technique (see Material and Methods) to find their parameters. This optimization yielded the Michaelis-Menten dependencies with the *K*_m_ values of 15, 161, and 409 μM (Fig. 1b, insert) whose linear combinations approximate the first, the second, and the third PC vectors with the square correlation coefficients (*R*_2_) of 0.9996, 0.9998, and 0.9507. The approximations of the first three PCs with linear combinations of these three components are shown in Fig. 1B main panel with lines.

**Figure 1.**
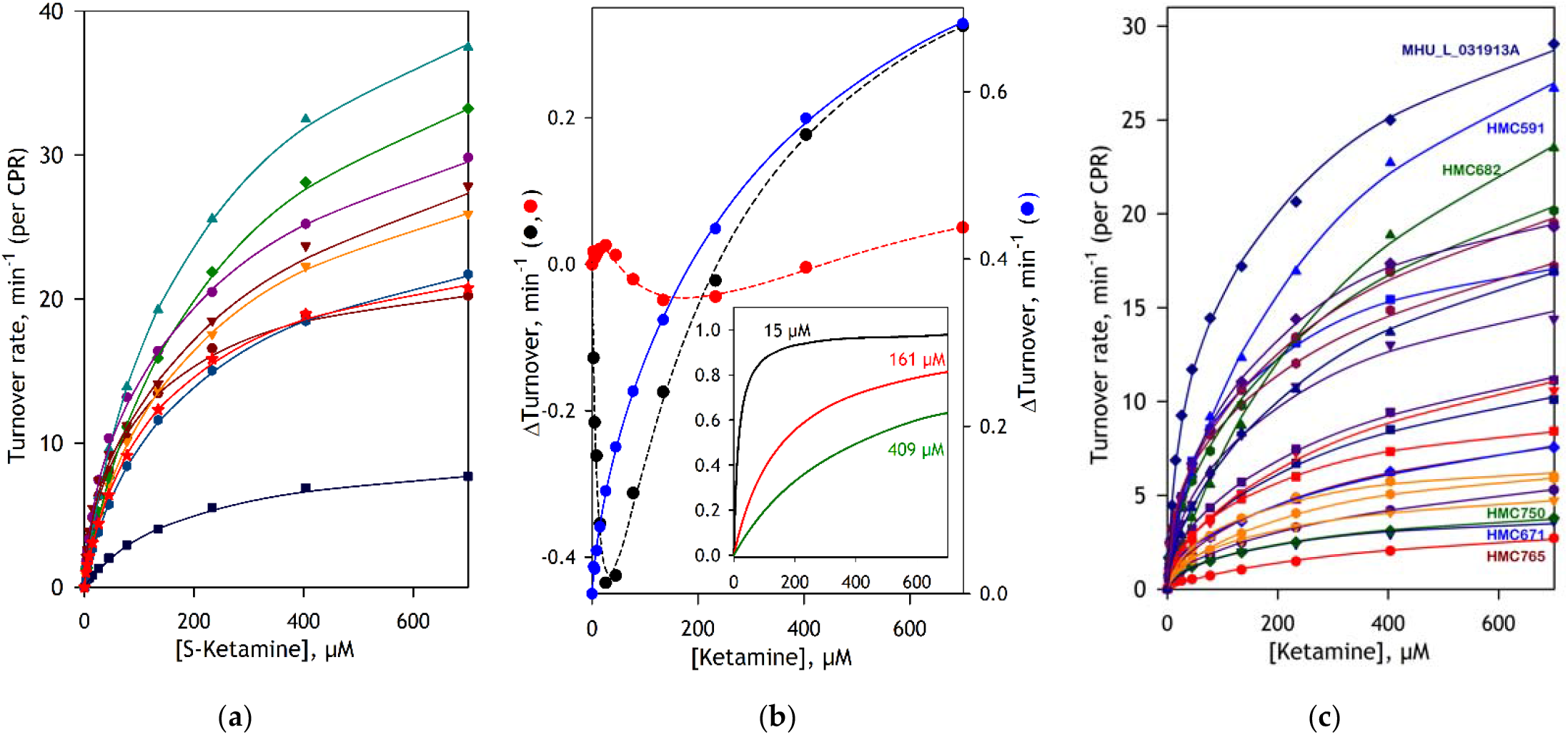
Substrate saturation profiles of N-demethylation of S-ketamine by HLM and their analysis by PCA. Panel a shows a dataset obtained with a set of 9 pooled HLM preparations. The solid lines represent the approximation of the SSPs by linear combinations of three Michaelis-Menten dependencies with globally optimized *K*_m_ values. The results of the application of PCA to this dataset are shown in panel b. The vectors of the first, second, and third principal components are shown in the main panel in blue, black, and red, respectively. The solid and dashed lines in the main panel show their approximations by linear combinations of three Michaelis-Menten dependencies with globally optimized *K*_m_ values, which are shown in the insert. The set of SSPs obtained with 23 individual SSP samples along with their approximations is exemplified in the panel c. Each SSP shown in panels a and c represents an average of the results of 3 - 6 individual experiments.

A combination of the multiplication coefficients obtained from this fitting with the respective vectors of eigenvalues was used to determine the amplitude of each of the three Michaelis-Menten components for each of the probed HLM samples and obtain the respective fitting traces, which approximate the experimental datasets with *R*_2_0.998 (Fig. 1a, solid lines). According to the results of this fitting, the first Michaelis-Menten component (*K*_m_=15 μM), which presumably reflects the participation of CYP2C19 and CYP4A11, constitutes from 1 to 21% (9.6% on average) of the total. The amplitude of the second component (*K*_m_ =161μM), which may be attributed to the involvement of the CYP3A enzymes, is highly variable and accounts for 0 - 81% of the total (average of 46.5%). The third component (*K*_m_ =409 μM), which apparently reflects the activity of CYP2C9, CYP1A2, CYP2D6, and CYP2E1, varies from 0 to 82% in the amplitude (43.9% on average). These results do not comply well with the expectations from the proportional projection approach (Table 1). Thus, the averaged amplitude of the low-affinity component (*K*_m_ =409 μM) is considerably higher than the predicted involvement of the low-affinity enzymes (CYP2C9, CYP1A2, CYP2D6, and CYP2E1). The amplitude of the high-affinity component (*K*_m_=15 μM) is also too high compared to expectations for the involvement of CYP2C19 and CYP4A11 (Table 1).

The set of SSPs for N-demethylation of ketamine obtained with this series of 23 HLM samples from individual donors is shown in Fig. 1c. The solid lines in this figure represent the results of global fitting of the dataset with a combination of three Michaelis-Menten equations with the *K*_m_ values of 15, 161 and 409 μM determined from the fitting of the pooled HLM dataset as described above. The square correlation coefficients for the fitting of the individual SSPs were always >0.990 and, in most cases, exceeded 0.997. As seen from this figure, different HLM samples differ by up to an order of magnitude in the amplitude of SSPs. The shape of the titration curves also varies dramatically. The range of variation of the relative amplitude of the high-affinity component (*K*_m_=15 μM) was from 0 to 29%, which is similar to that observed for the pooled HLM preparations. However, the distribution of the amplitudes between the two lower affinity phases differed considerably from that observed in the pooled samples. Here the averaged amplitude of the component with *K*_m_=161 μM was equal to 25%, which is notably lower than that characteristic of pooled preparations (46.5%). The fraction of the lowest affinity component varied from 0 to 99% with an average of 63.5%. One of the possible reasons for this difference may be a large fraction of heavy drinkers (7 out of 23) in the list of HLM donors.

To probe the effect of alcohol consumption on ketamine metabolism we assessed the dependence of the overall turnover rate (*V*_max total_) and the distribution between the three individual kinetic components of the ketamine metabolism on the Provisional Index of Alcohol Exposure (PIAE) introduced in our recent study of the effect of alcohol exposure on the proteome of the drug-metabolizing enzymes and transporters in the human liver [27]. PIAE is the combination of the relative abundances of four protein markers of alcohol exposure the heat-shock 70 protein HSPA5 (GRP75), protein disulfide isomerases P4HB and PDIA3, and carboxylesterase CES2. A strict correlation of PIAE with the level of alcohol consumption reported by liver donors has been demonstrated in our previous study [27]. This index varies from ∼0 to ∼5, and its numeric value roughly approximates the average number of alcoholic drinks per day consumed by the liver donor.

The dependency of *V*_max total_ on PIAE is shown in Fig. 2a. As seen from this figure, the overall rate of ketamine demethylation exhibits a pronounced increase at increasing PIAE. Excluding two outliers with the highest PIAE values (liver samples HMC593 and FHU-L-050211), its proportionality to PIAE is characterized by the correlation coefficient of 0.48, which corresponds to the Student’s T-test *p*-value of 0.03. Importantly, the exclusion of the HLM samples of homo- and heterozygotes of CYP2B6*5 and CYP2B6*6 from consideration improves the correlation and results in T-test *p*-value as low as 0.012 (Fig. 2A, dashed line). In contrast, no considerable effect of CYP2D6 SNPs was observed.

**Fig 2.**
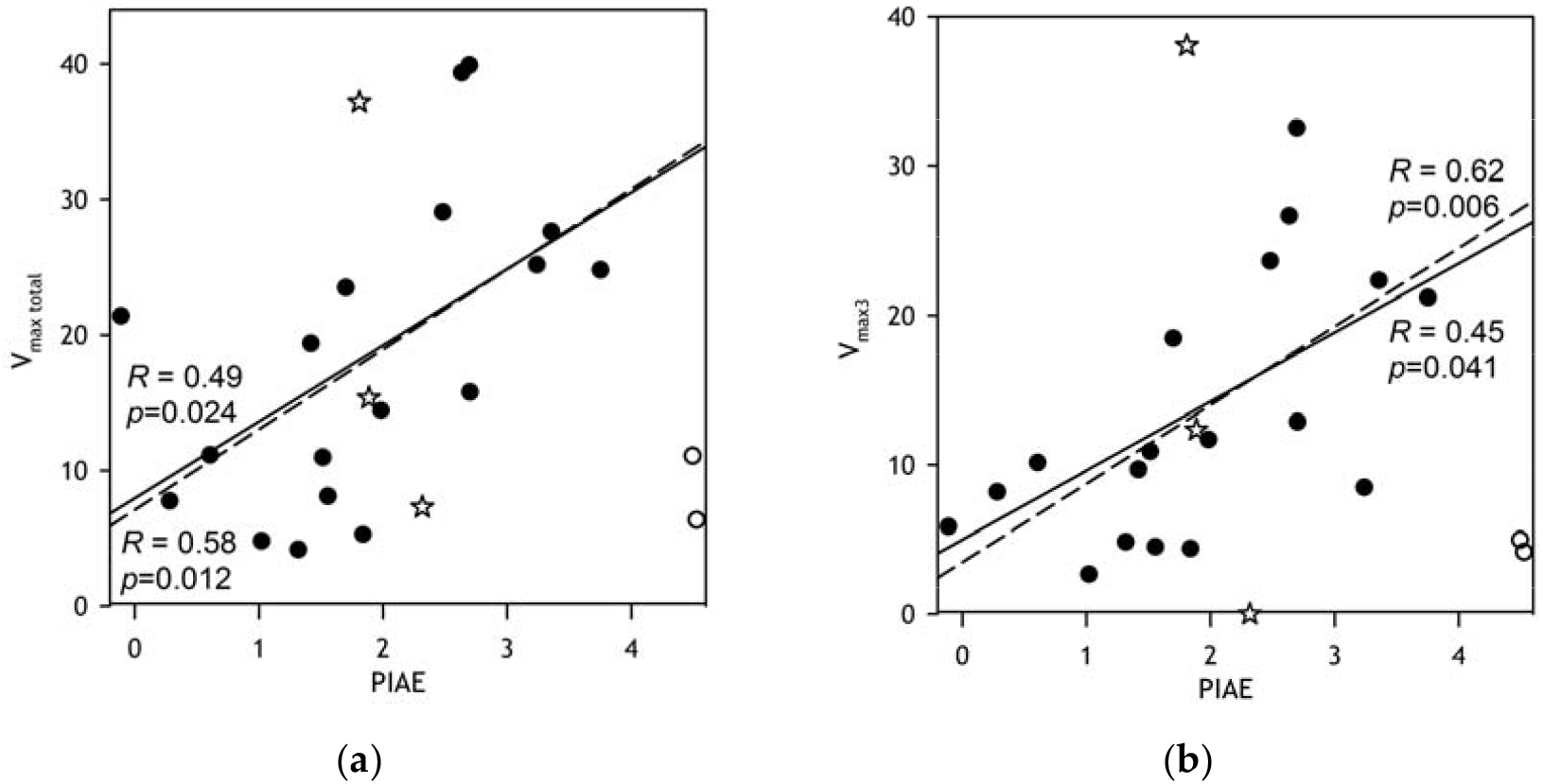
Correlations between the total maximal rate (*V*_max total_, panel a) and the maximal rate of the low-affinity component (*V*_max3_, panel b) of ketamine demethylation with the provisional index of alcohol exposure (PIAE) of liver donors. The solid line shows the linear approximation of the whole dataset excluding two apparent outliers at very high PIAE shown with open circles. The dashed line shows the linear approximation of the dataset with the points corresponding to homo- and heterozygotes of rare CYP2B6 variants (open stars) excluded.

As illustrated in Fig. 2b, the increase in V_max total_ upon increasing PIAE is mainly due to the increasing amplitude of the low-affinity component (*V*_max3_). Linear approximation of the dependence of V_max3_ on PIAE with CYP2B6*5 and CYP2B6*6 homo- and heterozygotes excluded (Fig. 2B, dashed line) yields the correlation coefficient of 0.62 (*p*=0.006). In contrast, no statistically significant correlation of *V*_max1_ and *V*_max2_ with the level of alcohol exposure was observed (data not shown).

Based on the linear approximation shown in the dashed line in Fig. 2a, we can estimate that in the heavy drinkers (PIAE=3.5) the maximal rate of ketamine metabolism in the liver is approximately five times higher than in non-drinkers (PIAE=0.5).

### 3.4 Metabolism of amitriptyline by HLM

A series of titration curves of amitriptyline demethylation obtained with a series of nine pooled HLM preparations is shown in Fig. 3a. Similar to the case of ketamine, the SSPs obtained with different pooled HLM preparation demonstrate high variability in their shape and amplitude. The application of PCA for the global fitting of this dataset is illustrated in Fig. 3B. Here, the best approximation of the first three principal components was obtained with a combination of three Michaelis-Menten dependencies with *K*_m_ values of 5, 160, and 671 μM. Their linear combinations approximate the first, the second, and the third PC vectors with the square correlation coefficients (*R*_2_) of 0.9987, 0.9917, and 0.8910 (Fig. 3B, main panel, solid lines).

**Fig 3.**
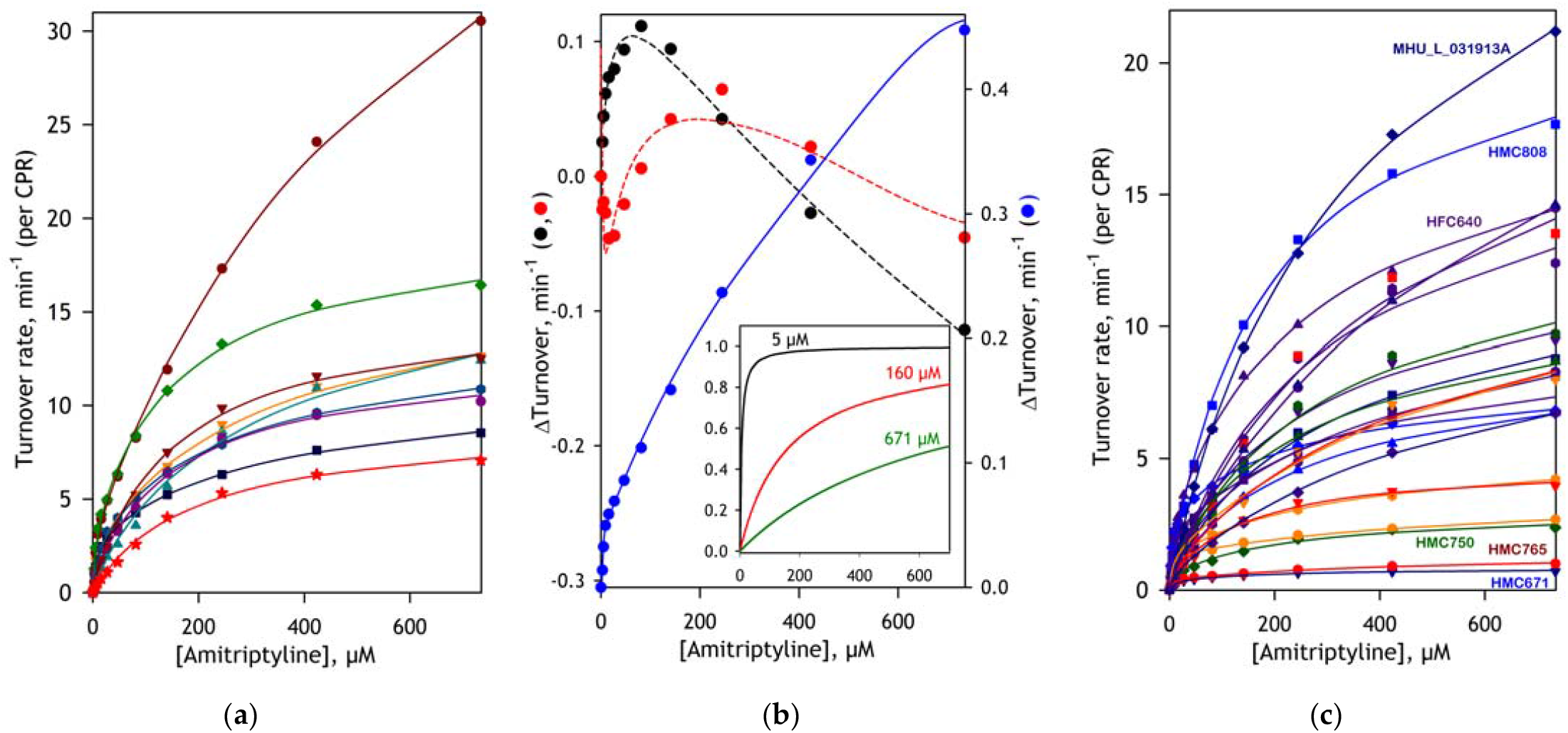
Substrate saturation profiles of N-demethylation of amitriptyline by HLM and their analysis by PCA. Panel a shows a dataset obtained with a set of nine pooled HLM preparations. The solid lines represent the approximation of the SSPs by linear combinations of three Michaelis-Menten dependencies with globally optimized *K*_m_ values. The results of the application of PCA to this dataset are shown in panel b. The vectors of the first, second, and third principal components are shown in the main panel in blue, black, and red, respectively. The solid and dashed lines in the main panel show their approximations by linear combinations of three Michaelis-Menten dependencies with globally optimized *K*_m_ values, which are shown in the insert. The set of SSPs obtained with 23 individual SSP samples along with their approximations is exemplified in the panel c. Each SSP shown in panels a and c represents an average of the results of 3 - 6 individual experiments.

Results of the approximation of the dataset shown in Fig. 3A with the combination of these three Michaelis-Menten dependencies (*R*_2_0.994) are shown in Fig. 3a in solid lines. Notably, the *K*_m_ value of the highest affinity component found in the global fitting (5 μM) is considerably lower than the *K*_m_ values characteristic for any of the amitriptyline-metabolizing P450 species found in our studies with recombinant enzymes (Table 2). According to the results of the global fitting, this Michaelis-Menten component constitutes 0 to 21% (8.6% on average) of the total SSP amplitude. It is also remarkable that the *K*_m_ values characteristic for CYP3A4 and CYP1A2 (69 and 56 μM, respectively), which are expected to be among the main amitriptyline metabolizers (see Table 2), do not find a close match in the set of three *K*_m_ values obtained in the global fitting. Presumably, the participation of these enzymes is reflected in the second Michaelis-Menten component (*K*_m_ =160 μM), along with the activity of CYP2C9 and CYP2E1 (259 and 205 μM, respectively). The fractional content of this component varies from 0 to 97%. The *K*_m_ value of the third, lowest affinity, component (671 μM) also finds no match in the set of kinetic parameters of 11 P450 enzymes probed for amitriptyline metabolism. This component constitutes from 0 to 94% of the total SSP amplitude.

The set of SSPs for N-demethylation of amitriptyline obtained with our set of 23 individual HLM samples is shown in Fig. 3c. The solid lines in this figure represent the results of global fitting of the dataset with a combination of three Michaelis-Menten equations with the *K*_m_ values of 5, 160 and 671 μM determined above with the pooled HLM dataset. The square correlation coefficients for this fitting were >0.99 and, in most cases, exceeded 0.995. Similar to the case of ketamine (Fig. 1), the amplitude and the shape of SSPs of the individual HLM samples varied dramatically. The fraction of the highest affinity component (*K*_m_ =5 μM) varied from 0 to 29% (10% on average), while the fractions of the 160 μM and 671 μM components varied from 0 to 90% and from 0 to 95% with the averages of 42 at 48%, respectively.

Investigating the relationship between the amplitudes of the three Michaelis-Menten components and the level of alcohol exposure of the donors, in contrast to the case of ketamine, we did not detect a significant correlation of *V*_max total_ with the PIAE. Instead, we detected a pronounced increase in the fraction of the third, low-affinity phase (*F*_3_) with increasing levels of alcohol consumption. As illustrated in Fig. 4a, the proportionality between *F*_3_ and PIAE in the full set of 23 HLM samples is characterized by the correlation coefficient of 0.39 and T-test p-value of 0.066 (Fig. 4). Although the *p*-value for this correlation is above the 0.05 threshold of statistical significance, the omission of the HLM samples of homo- and heterozygotes of rare variants of CYP2B6 and CYP2D6 boosts the correlation coefficient to 0.77 and decreases the *p*-value to 0.001 (Fig 4a, dashed line). As shown in Fig. 4b, this increase in the fraction of the low-affinity component at increasing PIAE takes place at the expense of the decreasing fraction of the middle-affinity component. We may conclude therefore that increased alcohol exposure reduces the role of medium-affinity P450 enzymes, and particularly CYP3A4, in the amitriptyline metabolism concomitant with increasing participation of the enzymes with low affinity to the substrate.

**Fig 4.**
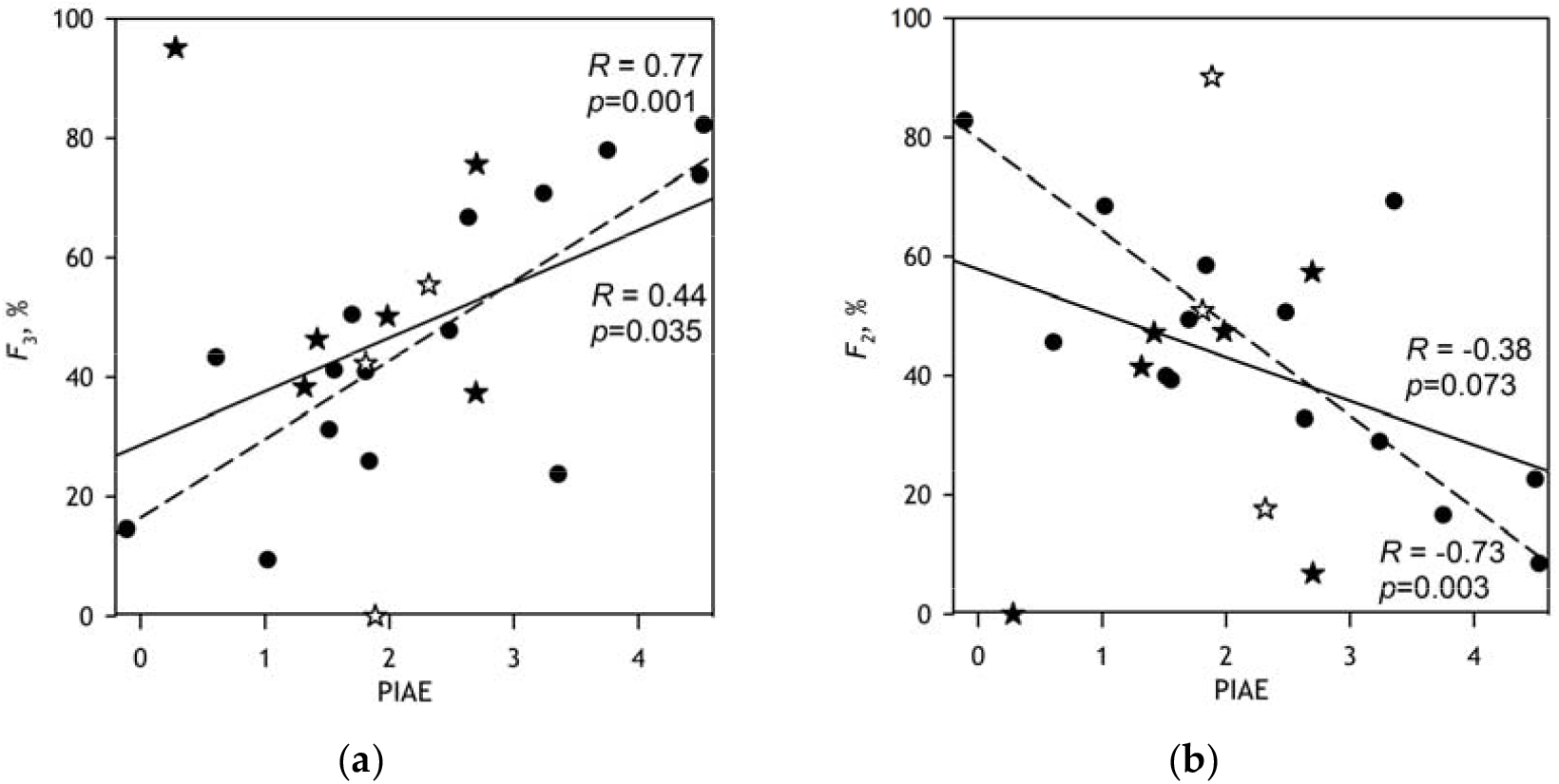
Correlations of the fractional amplitude of the low affinity (*F*_3_, panel a) and the medium affinity (*F*_2_, panel b) components of amitriptyline demethylation with the provisional index of alcohol exposure (PIAE) of liver donors. The solid lines show the linear approximation of the whole datasets and the dashed lines represent the approximation of the dataset with the points corresponding to homo and heterozygotes of rare variants of CYP2B6 (open stars) and CYP2D6 (closed stars) excluded.

### 3.5 Inhibition of N-demethylation of ketamine and amitriptyline by CYP3cide

To probe the role of CYP3A4, which, according to the proportional projection calculations (Tables 1 and 2), is expected to be the principal metabolizer of ketamine and amitriptyline, we studied the inhibition of the metabolism of both substrates by CYP3cide, a potent and highly specific inhibitor of CYP3A4 [50]. We probed the effect of CYP3cide on ketamine and amitriptyline metabolism in a series of 8 pooled HLM preparations. In all cases the inhibition was well-pronounced. In the case of ketamine, its amplitude comprises 55.5±2.5% and the *IC*_50_ amounts to 0.142±0.22 μM. The inhibition of amitriptyline metabolism by CYP3cide has an *IC*_50_ value of 0.065±0.15 μM and varies in amplitude from 61 to 80% (average of 71±5%). These results are consistent with the primary role of CYP3A4 in the metabolism of both substrates in agreement with the proportional projection analysis (Tables 1 and 2).

Importantly, when analyzing the relationship between the amplitude of inhibition of ketamine demethylation by CYP3cide and the kinetic parameters of its metabolism by different pooled HLM preparations, we observed a significant correlation between the *V*_max_ of the high-affinity (*K*_m_ =5 μM) component and the depth of inhibition. This relationship is illustrated in Fig. 5A. It is characterized by a correlation coefficient of 0.735 and a T-test *p*-value of 0.038. This observation may signify a modulatory effect of CYP3A4 interactions on an enzyme responsible for the high-affinity component of ketamine metabolism, such as CYP2C19 or CYP4A11.

**Fig 5.**
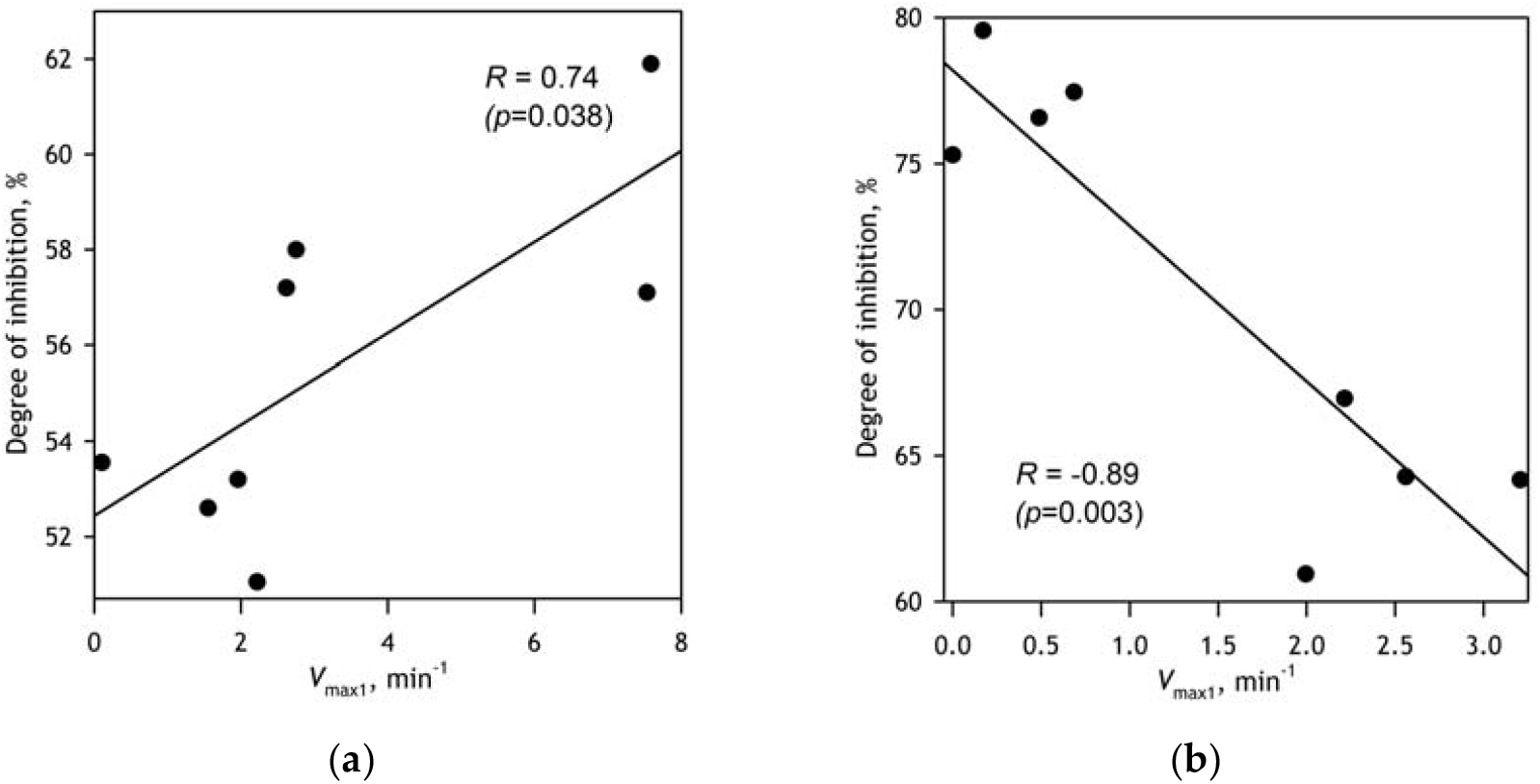
Correlations between the maximal rate of the high-affinity component of N-demethylation of ketamine (a) and amitriptyline (b) with the depth of inhibition of these reactions by CYP3cide.

In the case of amitriptyline, we detected a significant correlation of the opposite sign (Fig. 5B). Here the relationship between the depth of inhibition and *V*_max1_ is characterized by a correlation coefficient of -0.891 and the T-test *p*-value of 0.003.

### 3.6 Analysis of correlations between the composition of the cytochrome P450 ensemble and the kinetics of ketamine and amitriptyline demethylation

To further explore the role of the individual P450 species in the metabolism of ketamine and amitriptyline, we investigated the relationship between the composition of the P450 ensemble in HLM preparations and the shape and amplitude of SSPs of these substrates. In these studies we attempted to approximate the vectors of amplitudes of each of the three Michaelis-Menten components (*V*_max1_, *V*_max2_, and *V*_max3_) and their total (*V*_max total_) win linear combinations with up to four vectors of relative fractional content (VFC) of eleven drug-metabolizing cytochromes P450 (CYP1A2, CYP2A6, CYP2B6, CYP2C8, CYP2C9, CYP2C19, CYP2D6, CYP2E1, CYP3A4, CYP3A5, and CYP4A11). To do so we utilized the multidimensional linear least-square regression algorithm as described in [32]. This algorithm was sequentially applied to every possible combination of the VFCs of 1 - 4 species in the above set of P450 proteins. The approximation of the profile of *V*_max total_ of ketamine metabolism by 23 individual HLM samples by a combination of four VFCs is exemplified in Fig. 6.

**Fig 6.**
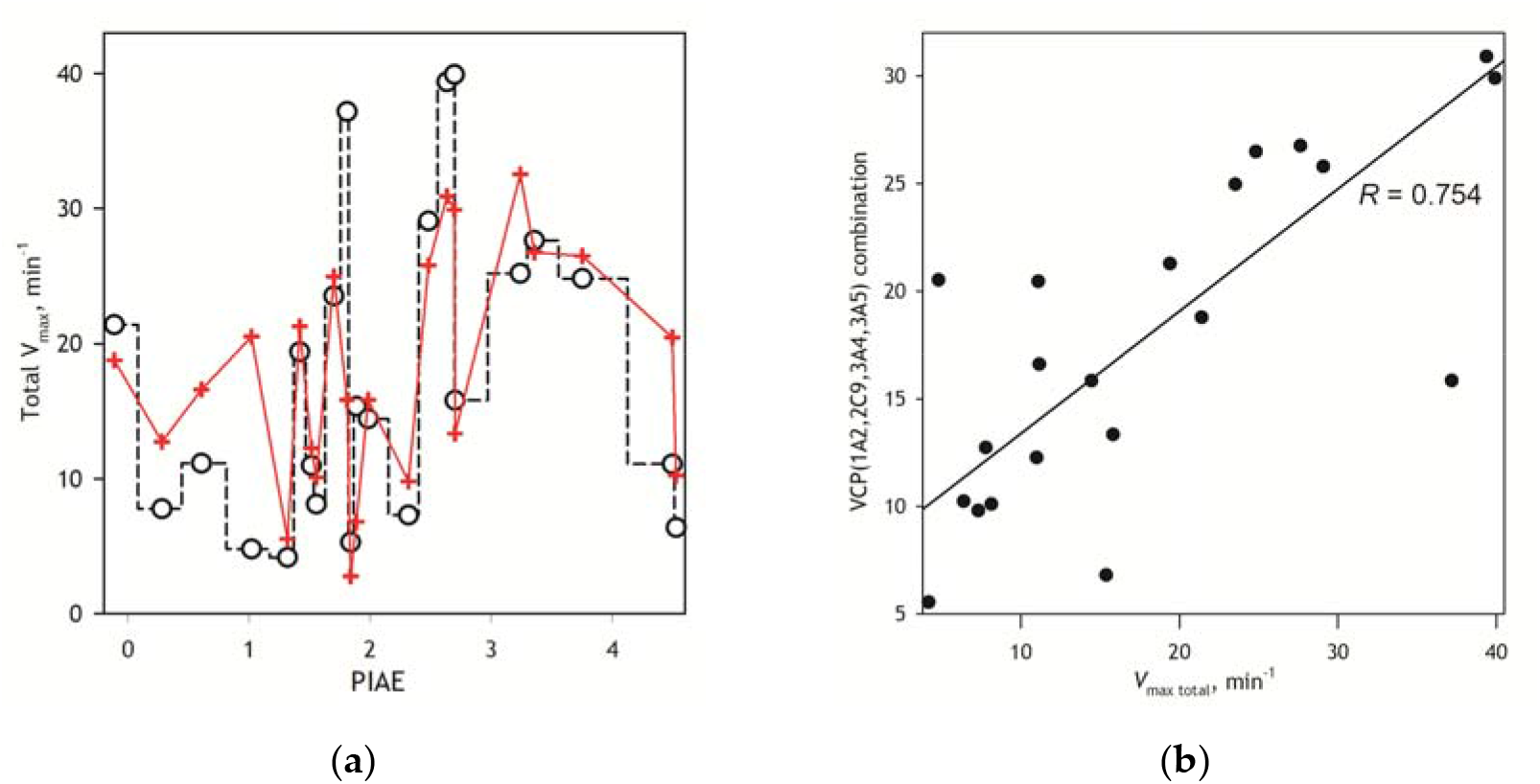
Analysis of correlations of the total maximal rate of ketamine demethylation with the composition of the cytochrome P450 pool in 23 HLM preparations from individual donors. Panel a shows the approximation of the dataset of *V*_max total_ values (black open circles) with a linear combination of the vectors of relative fractional content (VFC) of CYP1A2, CYP2C9, CYP3A4, and CYP3A4 taken with the multiplication factors of -9.3,-9, +7.2 and -3.7, respectively. The X-axis in this panel corresponds to the PIAE of the liver donors. Panel (b) shows the same approximation as a plot of the found VFC combination versus *V*_max total_.

We applied this procedure to the dataset obtained with 23 individual HLM samples and the combined dataset of 32 individual and nine pooled preparations. We also analyzed the dataset of individual 14 HLM samples, where the preparations from donors homo- or heterozygous for rare CYP2B6 and CYP2D6 variants were omitted. The results of these trials are summarized in Fig.7, which illustrates the process of finding the best combinations of four vectors of relative abundance of P450 species that approximate the profiles of the maximal turnover rate (*V*_max total_) and the amplitudes of each of the three individual Michaelis-Menten components (*V*_max1_, *V*_max2,_ and *V*_max3_) of the SSPs. In this analysis, the number of individual P450 profiles in approximation was increasing stepwise from 1 to 4. The numbers shown in the tables indicate the stage at which each P450 enzyme appeared in the approximating combination. The enzymes with a positive contribution are indicated by a red background, while the green background designates the proteins whose contribution is negative (i.e. inhibiting). The pink and light green shadowing indicates the species that appeared in the combination at the early steps but were replaced by other P450 species at the following stages.

As seen from Fig. 7a, in good agreement with the expectations, the maximal rate of ketamine metabolism reveals a pronounced correlation with the fractional content of CYP3A4. However, this is where the correspondence with the prediction ends. Not one of the other three predicted principal players in ketamine metabolism - CYP2B6, CYP2C9, and CYP1A2 - reveals a reproducible positive correlation between its relative fractional content and *V*_max total_. Right the opposite - the negative correlation observed with CYP1A2 and CYP2C9 suggests their inhibitory effects. A negative correlation is also observed with the fractional content of CYP3A5, which is best revealed in its effect on the low-affinity component.

**Fig 7.**
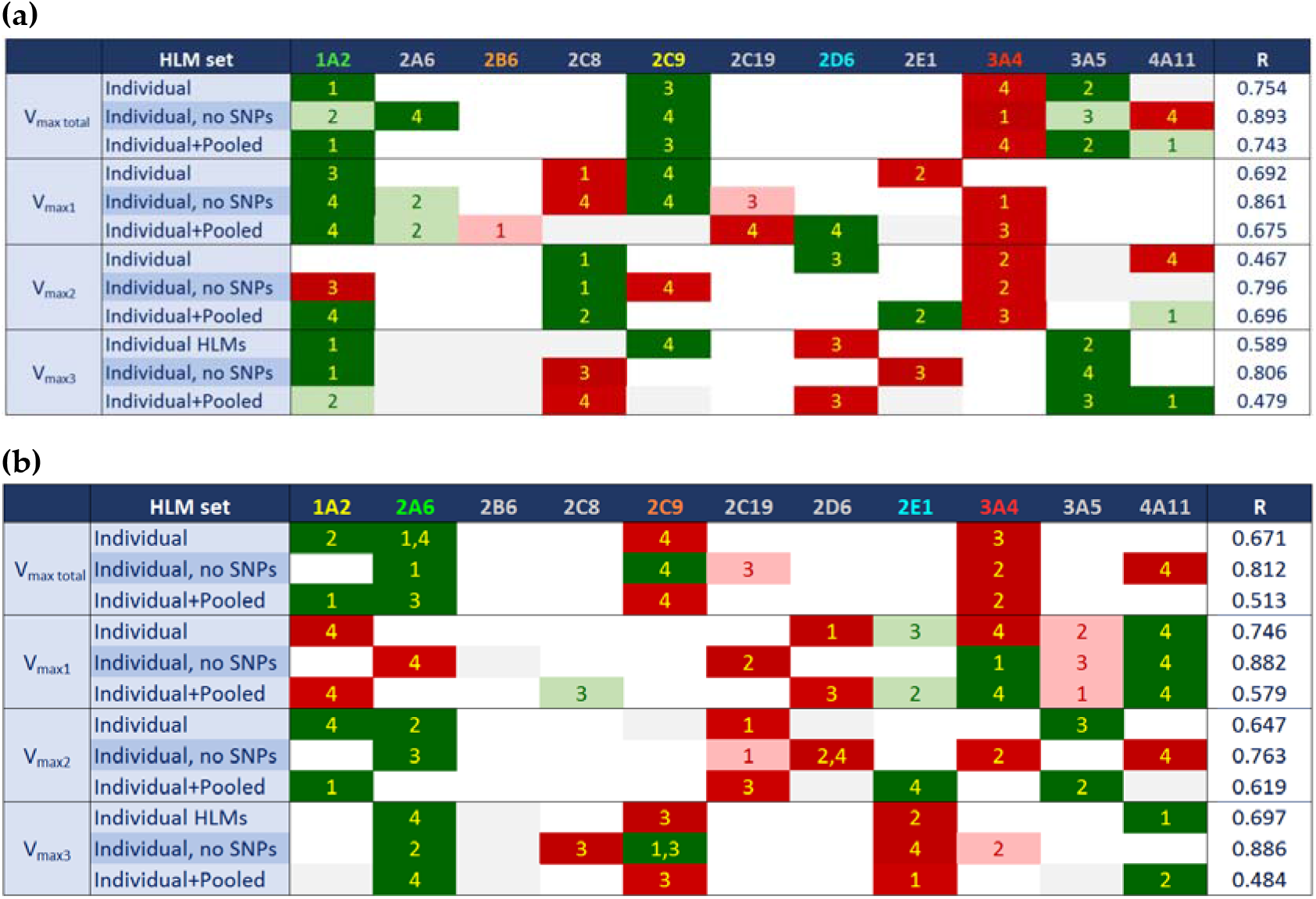
Correlations of Substrate Saturation Profiles of ketamine (A) and amitriptyline (B) demethylation with the composition of the cytochrome P450 pool in a set of 23 individual and 9 pooled HLM preparations. The tables illustrate the process of finding the best combinations of four profiles of relative fractional content of 11 major P450 species that approximate the profiles of *V*_max total_, *V*_max1_, *V*_max2,_ and *V*_max3_. The numbers shown in the table indicate the stage of successive approximations with 1 to 4 VFC combinations. The species with positive correlations are shown with red shadowing. The negative correlation is indicated by a green background. The light red and light green shadowing indicates the species that appeared in the early steps of the analysis but were replaced by other P450s later. The values in the rightmost column are the correlation coefficients of the four-component approximations. In the header rows, the P450 species predicted to play the major role in the metabolism are highlighted in red, orange, yellow, green, and blue colors where the red color designates the predicted primary metabolizer.

Analyzing the correlations with the *V*_max_ values of three individual Michaelis-Menten components (*V*_max1_, *V*_max2,_ and *V*_max3_), we see a positive correlation of CYP3A4 abundance with the amplitudes of the first (*K*_m_=15 μM) and the second (*K*_m_=161 μM) components. Correlation with the amplitude of the high-affinity component, where CYP3A4 (*K*_m_=113 μM, Table 1) is unlikely to be involved, presumably suggesting its activating effect on P450 enzyme(s) with high affinity to ketamine. This observation is in good agreement with a positive correlation between *V*_max1_ and the depth of inhibition of ketamine demethylation by CYP3cide (Fig. 5a) discussed above. Furthermore, concomitant with the expectations, the amplitude of the low-affinity phase exhibited a strong correlation with the abundance of CYP2D6, the most efficient low-affinity metabolizer.

The apparent inhibitory effect of CYP1A2 is best pronounced in the high-affinity component of SSP, while the inhibition by CYP3A5 is revealed for the lowest affinity (*K*_m_=409 μM) phase. Besides the effects of CYP1A2 and CYP3A5, our analysis suggests a negative correlation of the content of CYP2C8 with the amplitude of the middle affinity (*K*_m_=161 μM) component and CYP4A11 on the low-affinity phase.

In the case of amitriptyline, the correlations of *V*_max total_ with the abundances of CYP3A4 is also well pronounced. The correlation with the content of CYP2C9 is less evident. In the case of the other two predicted potent metabolizing enzymes, CYP1A2 and CYP2D6, we observed a strong negative correlation suggesting their inhibitory effect.

Another strong relationship revealed by our analysis is a positive correlation of *V*_max3_ with CYP2E1 content, which agrees with the important role of this enzyme in the low-affinity metabolism of amitriptyline suggested by proportional projection analysis. On the flip side, the strong positive correlation of the amplitude of the high-affinity component (Km = 5 μM) with the fractional content of CYP1A2 contradicts the relatively low affinity (Km = 56 μM) characteristic of the recombinant CYP1A2 enzyme. Likewise, the apparent increase in *V*_max2_ (*K*m=160 μM) with increasing CYP2C19 abundance is not consistent with its high affinity for amitriptyline (Km=14 μM).

In summary, while confirming the important role of CYP3A4 in the metabolism of both substrates and corroborating the involvement of CYP2D6 and CYP2E1 in the low-affinity components of SSPs of ketamine and amitriptyline, respectively, our correlational analysis failed to substantiate other inferences deduced by the proportional projection approach. Instead, it demonstrated a very complex mutual functional effects of multiple P450 species.

## 4. Discussion

In this study, we used a combination of the high-throughput kinetic assays with global proteomic analysis to explore the correlations of the kinetic profiles of ketamine and amitriptyline metabolism with the composition of the P450 ensemble in HLM and the level of alcohol exposure of liver donors. One of our aims was to probe the applicability of the proportional projection (total normalized rate) approach for predicting the pathways of metabolism of drugs metabolized by multiple cytochrome P450 enzymes. We sought to probe the instances of non-additivity of the functional properties of the individual members of the microsomal cytochrome P450 ensemble and identify possible deviations of the proportional projection rule. Another goal was to probe the effects of chronic alcohol exposure on the function of the cytochrome P450 ensemble and analyze their linkages with the alcohol-induced changes in the composition of the P450 pool.

The results of our recent study of HLM proteome in a set of 96 HLM samples from donors with different levels of alcohol consumption [27] allowed us to establish a biomarker-based provisional index of alcohol exposure (PIAE). This parameter was used here to explore the impact of alcohol exposure on the metabolism of ketamine and amitriptyline, the drugs used as antidepressants in patients with alcohol withdrawal syndrome.

Studying the correlations between the SSPs of ketamine demethylation and PIAE, we evidenced an important increase in the maximal rate of ketamine demethylation by chronic alcohol consumption. This effect is mainly due to an increase in the *V*_max_ of the low-affinity component of the reaction. Among the possible reasons for this effect may be an alcohol-induced increase in the abundance of CYP2E1, which is involved in the low-affinity ketamine metabolism along with CYP2D6. Another possible factor causing an increasing *V*_max total_ in alcohol consumers may be an alcohol-induced increase in the abundance of CPR, which we demonstrated in our proteomic study [27]. However, even considered together, these changes in protein expression are unlikely to be capable of causing such an important increase in *V*_max total_ (up to 5 times). Thus, the mechanism of alcohol-induced boost in ketamine metabolism requires further exploration.

The analysis of correlations between the kinetics of amitriptyline demethylation and PIAE also demonstrates an important increase in the role of low-affinity enzymes by alcohol exposure. However, in this case, this effect does not affect the *V*_max total_ as it takes place at the expense of the decreasing fraction of the middle-affinity component. We may hypothesize therefore that increased alcohol exposure reduces the role of medium-affinity P450 enzymes, and particularly CYP3A4, in the amitriptyline metabolism concomitant with increasing participation of enzymes responsible for its low-affinity metabolism.

Remarkable complexity in the mechanisms of modulation of drug metabolism by alcohol exposure is also revealed in the observed impact of rare genetic variants of CYP2B6 and CYP2D6 on correlations between PIAE and the kinetic parameters of metabolism of ketamine and amitriptyline. Thus, in the case of ketamine, the exclusion of homo- and heterozygotes of minor genetic variants of CYP2B6, a medium-affinity metabolizer (*K*_m_=54μM), from the analysis considerably improved the correlation between the degree of alcohol exposure and the amplitude of the low-affinity (Km=409 μM) phase. This observation is difficult to explain by the direct involvement of CYP2B6 in ketamine metabolism. Instead, it may reveal the effects of CYP2B6 on the activity of the low-affinity enzymes. Similarly, indirect effects of interactions between P450 enzymes and/or their competition for CPR are apparently revealed in an essential increase in correlation of PIAE with the distribution between the medium- and low-affinity phases upon omission of minor variants of CYP2B6 and CYP2D6 (Fig. 4), which are both characterized with high affinity to amitriptyline (see Table 2).

Besides studying the effects of alcohol consumption on drug metabolism, this study is aimed at the exploration of functional non-additivity in the drug-metabolizing ensemble of the human liver [5]. It further builds on the use of a combination of proteomic profiling of drug-metabolizing enzymes with a global analysis of saturation profiles of polyspecific drug substrates introduced in our previous publication [22] as an instrument for these studies. Here we elaborated a new, improved version of our initial method where the PCA transform is used for a global fitting of a set of SSPs with a linear combination of several Michaelis-Menten equations with globally optimized *K*_m_ values. The resulting sets of amplitudes of the individual (high-, medium- and low-affinity) components allows more straightforward interpretation than the sets of PCA eigenvalues used in our initial study. To our knowledge, this is the first attempt to use global kinetic analysis of a set of substrate saturation profiles to characterize the variability of drug metabolism in a series of individual HLM samples. Our new global fitting strategy allowed us to probe the involvement of P450 enzymes with different affinities to a substrate in its metabolism in each HLM preparation. The knowledge of the differences between HLM samples in the composition of the P450 pool deduced from the proteomic analysis was then used to examine the correlations between the kinetic profiles between the fractional content of eleven major P450 enzymes.

Although some of the conclusions from our correlational analysis comply with the initial predictions, the integral picture of correlations reveals critical limitations of the proportional projection approach and demonstrates a pronounced functional non-additivity in the P450 ensemble. Thus, neither CYP1A2, CYP2B6 nor CYP2C9, which were predicted to be among the principal metabolizers of ketamine, exhibit a pronounced positive correlation with the rate of its metabolism. Instead, the inverse was found - the abundances of CYP1A2, CYP2C9, and CYP3A5 were negatively correlated with the total rate of ketamine metabolism (see Fig. 7A) suggesting their inhibitory effects. Furthermore, CYP2A6, which is predicted to play an important role in the high-affinity metabolism of amitriptyline, demonstrated a negative correlation of its fractional content with the rate of its demethylation, which is best revealed in the amplitude of the low-affinity component.

In total, these observations demonstrate a lack of straight additivity in the functional properties of the members of the microsomal P450 ensemble. They reveal a network of complex mutual functional effects of multiple P450 species that largely limits the applicability of the proportional projection approach to predicting drug metabolism.

These results are in good agreement with the concept of functional integration of multiple P450 species via their protein-protein interactions and the formation of heteromeric P450 complexes in the microsomal membrane [4-5,29,51]. According to this concept, a significant part of the cytochrome P450 pool in HLM is deposited in the inactive (“latent”) positions in heteromeric complexes of multiple P450 species. Availability of a particular cytochrome P450 for interaction with substrates and reductase and, thus, its involvement in catalytic activity is presumed to be a complex function of the preferences of individual species for occupying the “active” and “latent” positions, the composition of the P450 pool and the presence of selective substrates of individual P450 species.

The approaches introduced in this study offer a potent strategy for in-depth exploration of the functional interactions between the major drug-metabolizing P450 species in their native environment. Further studies in this direction are vital for untangling the network of functional interactions between P450 enzymes and better understanding the interrelationships between the composition of the P450 ensemble and its potency to metabolize any particular substrate (its metabolic profile).

## Author Contributions

Dmitri Davydov: Conceptualization, Formal analysis, Funding acquisition, Investigation, Methodology, Software, Writing - original draft; Writing - review & editing; Bhagwat Prasad: Formal analysis, Funding acquisition, Methodology, Writing - review & editing; Phil- lip Lazarus: Methodology, Resources, Formal analysis, Writing - review & editing; Kannapiran Ponraj: Investigation, Formal Analysis, Writing - original draft; Kari A. Gaither: Investigation, Formal analysis; Dilip Kumar Singh: Investigation, Formal analysis; Nadezhda Davydova: Investigation; Mengqi Zhao: Investigation; Shaman Luo: Investigation.

## Acknowledgments

This research was supported by the National Institute on Alcohol Abuse and Alcoholism of the National Institutes of Health under Award R01AA030155.

